# Minimal Impact of Low Vision on Explicit Sensorimotor Adaptation

**DOI:** 10.1101/2025.07.29.667489

**Authors:** Mihai Cipleu, Sritej Padmanabhan, Emily A. Cooper, Jonathan S. Tsay

## Abstract

Rehabilitation from motor system dysfunction relies on learning deliberate motor corrections through practice and feedback. This is called explicit motor adaptation. One key source of feedback for this adaptation is the visual error signal between the intended movement and the achieved movement. As people age, both motor dysfunction and visual impairment become more common, potentially compromising the visual feedback signal. Previous work has shown that visual impairment can disrupt the implicit, automatic adjustments made by the sensorimotor system. But how visual impairment influences the explicit motor adaptation, a cornerstone of rehabilitation, remains unknown. To address this gap, we recruited individuals with low vision—defined as uncorrectable visual impairment resulting in functional vision loss—and age-matched controls to complete a visuomotor task designed to isolate two components of explicit motor adaptation: discovering a new deliberate sensorimotor strategy and recalling a previously learned one. Surprisingly, low vision had no measurable impact on either component. Despite reduced visual fidelity, individuals with low vision were as effective as controls in both discovering and retrieving successful explicit sensorimotor strategies. These results highlight potential mechanisms that can be leveraged in rehabilitation.

## Introduction

Sensorimotor adaptation—the process of correcting movement errors through feedback and practice— is a fundamental human capacity that keeps our actions well-calibrated amid changes to the body and environment (Shadmehr et al., 2010; Tsay, Kim, et al., 2024). Motor adaptation does not rely on a single mechanism but instead recruits multiple learning processes (Hadjiosif & Krakauer, 2021; Kim et al., 2020; Maresch et al., 2021; McDougle et al., 2016). A key distinction is between implicit adaptation (Kim et al., 2018; Mazzoni & Krakauer, 2006), which automatically recalibrates the sensorimotor system, and explicit strategies, which involve deliberate motor corrections to achieve task success (Hegele & Heuer, 2010; Seidler & Carson, 2017; Taylor et al., 2014). For instance, Stephen Curry may automatically compensate for arm fatigue to preserve shot stability (implicit adaptation) while deliberately alter his shot trajectory to avoid a defender (explicit strategy).

Extensive research has shown that feedback uncertainty weakens implicit adaptation. Consistent with Bayesian principles (Bays & Wolpert, 2007; Faisal et al., 2008; Körding & Wolpert, 2004; Trommershäuser et al., 2008; Tsay, Kim, et al., 2022), uncertain visual feedback reduces the reliability of the visual error signal—the mismatch between perceived and intended movement—thereby attenuating adaptation. This effect has been observed under degraded visual conditions (e.g., blur, low lighting) (Burge et al., 2008; Casasnovas et al., 2024; Crossley et al., 2025; Makino et al., 2023; Tsay, Avraham, et al., 2021; Wei & Körding, 2010; Z. Zhang et al., 2024) and in individuals with low vision— defined as uncorrectable visual impairment that results in functional vision loss—where uncertainty is intrinsic to the sensorimotor system (Tsay et al., 2023).

In contrast, the impact of visual uncertainty on explicit sensorimotor strategies remains unknown. This gap is surprising given that, like implicit adaptation, explicit strategies also heavily rely on visual error signals to correct errors in everyday tasks such as adjusting to an unfamiliar computer trackpad or compensating for wind conditions during outdoor sports. In explicit adaptation, these signals are essential for formulating and implementing deliberate motor corrections. While this learning should in principle be sensitive to degraded visual input, the extent of this sensitivity is unknown.

This lack of insight into how visual uncertainty affects explicit adaptation carries important clinical implications: Explicit adaptation is foundational to rehabilitation, where patients are often taught to deliberately compensate for sensorimotor deficits—such as muscle weakness or limb disuse— following neurological injury (Leech et al., 2021; Roemmich & Bastian, 2018). As people age, their probability of both sensorimotor dysfunction and visual impairment increase (Cisneros et al., 2024; Vandevoorde & Orban de Xivry, 2020; Varma et al., 2016; A. Zhang et al., 2025). As such, clarifying how visual uncertainty shapes strategy use is essential for advancing our mechanistic understanding of motor adaptation and for optimizing interventions that harness strategic compensation to help individuals with low vision regain motor control lost due to degraded central vision (e.g., age-related macular degeneration) and peripheral vision (e.g., glaucoma, retinitis pigmentosa).

To address this gap, we recruited individuals with low vision and age-matched controls to perform a visuomotor adaptation task designed to isolate two components of strategic adaptation: the discovery of a new strategy and the recall of a previously learned one (Brudner et al., 2016; Kitazawa et al., 1995; Tsay, Schuck, et al., 2022). The performance of both groups on both learning components was highly similar, suggesting that explicit adaptation can be robust to common levels of visual impairment. Together with previous work, these findings identify explicit adaptation as a viable mechanism to guide rehabilitative strategies for individuals with a combination of motor dysfunction and visual impairment.

## Results

How does low vision affect explicit adaptation? To answer this question, we tested participants with low vision (LV) and age-matched controls (N = 23/group) on a visuomotor rotation task designed to isolate the use of an explicit aiming strategy (Table 1). Participants used their computer trackpad to make rapid, goal-directed movements toward a visual target presented on the screen (Fig. 1A). After an initial veridical feedback baseline block to familiarize participants with the task environment and apparatus, the feedback cursor was rotated by 60° (Fig. 1B). To compensate for this rotation, both groups exhibited significant movement angle changes in the opposite direction of the rotation, drawing the cursor closer to the target (Fig. 1C). When participants were asked to forgo their strategies and re-aim back to the target during the initial no-feedback aftereffect block, both groups were able to “switch-off” their strategies and successfully re-aim back to the target. Both groups exhibited minimal aftereffects, confirming that our delayed feedback manipulation was successful in eliminating implicit adaptation (Brudner et al., 2016; Kitazawa et al., 1995). Upon re-exposure to the rotation in the Recall block, participants were able to rapidly retrieve their re-aiming strategy and subsequently switch back to aiming directly to the target in the second no-feedback aftereffect block. In sum, the adaptive changes observed in response to the 60° rotation were the result of strategic re-aiming, rather than implicit adaptation.

**Table 1.**
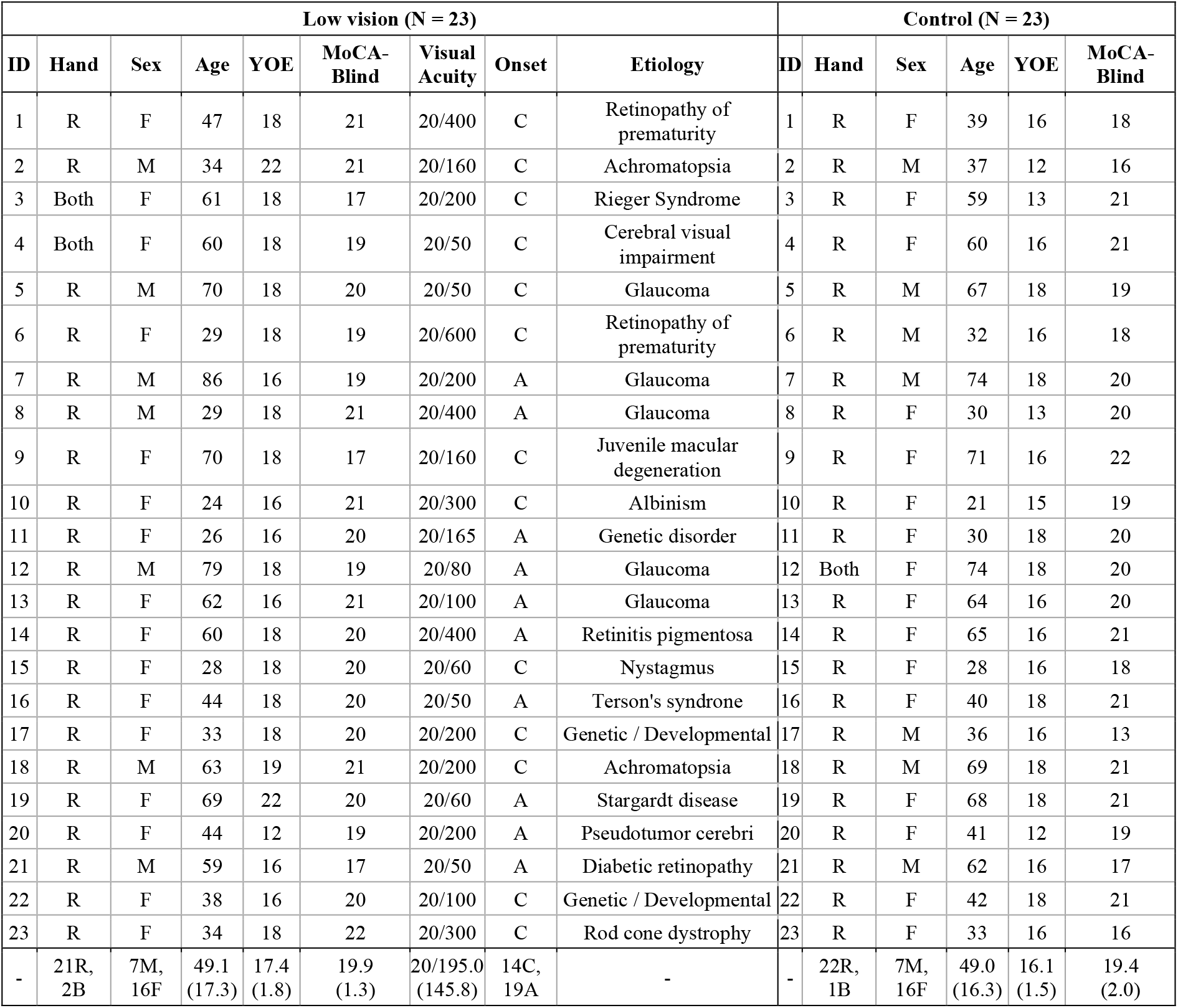
Demographic information for participants in the Low Vision and Control groups. Reported for each participant are ID, handedness, sex, age, years of education (YOE), visual acuity of their better-seeing eye, and MoCA-Blind scores. For participants with low vision, we also provide their self-reported etiology. Scores ≥ 18 on the MoCA-Blind indicate no cognitive impairment.

| Low vision (N = 23) |  |  |  |  |  |  |  |  | Control (N = 23) |  |  |  |  |  |
| --- | --- | --- | --- | --- | --- | --- | --- | --- | --- | --- | --- | --- | --- | --- |
| ID | Hand | Sex | Age | YOE | MoCA-Blind | Visual Acuity | Onset | Etiology | ID | Hand | Sex | Age | YOE | MoCA-Blind |
| 1 | R | F | 47 | 18 | 21 | 20/400 | C | Retinopathy of prematurity | 1 | R | F | 39 | 16 | 18 |
| 2 | R | M | 34 | 22 | 21 | 20/160 | C | Achromatopsia | 2 | R | M | 37 | 12 | 16 |
| 3 | Both | F | 61 | 18 | 17 | 20/200 | C | Rieger Syndrome | 3 | R | F | 59 | 13 | 21 |
| 4 | Both | F | 60 | 18 | 19 | 20/50 | C | Cerebral visual impairment | 4 | R | F | 60 | 16 | 21 |
| 5 | R | M | 70 | 18 | 20 | 20/50 | C | Glaucoma | 5 | R | M | 67 | 18 | 19 |
| 6 | R | F | 29 | 18 | 19 | 20/600 | C | Retinopathy of prematurity | 6 | R | M | 32 | 16 | 18 |
| 7 | R | M | 86 | 16 | 19 | 20/200 | A | Glaucoma | 7 | R | M | 74 | 18 | 20 |
| 8 | R | M | 29 | 18 | 21 | 20/400 | A | Glaucoma | 8 | R | F | 30 | 13 | 20 |
| 9 | R | F | 70 | 18 | 17 | 20/160 | C | Juvenile macular degeneration | 9 | R | F | 71 | 16 | 22 |
| 10 | R | F | 24 | 16 | 21 | 20/300 | C | Albinism | 10 | R | F | 21 | 15 | 19 |
| 11 | R | F | 26 | 16 | 20 | 20/165 | A | Genetic disorder | 11 | R | F | 30 | 18 | 20 |
| 12 | R | M | 79 | 18 | 19 | 20/80 | A | Glaucoma | 12 | Both | F | 74 | 18 | 20 |
| 13 | R | F | 62 | 16 | 21 | 20/100 | A | Glaucoma | 13 | R | F | 64 | 16 | 20 |
| 14 | R | F | 60 | 18 | 20 | 20/400 | A | Retinitis pigmentosa | 14 | R | F | 65 | 16 | 21 |
| 15 | R | F | 28 | 18 | 20 | 20/60 | C | Nystagmus | 15 | R | F | 28 | 16 | 18 |
| 16 | R | F | 44 | 18 | 20 | 20/50 | A | Terson's syndrome | 16 | R | F | 40 | 18 | 21 |
| 17 | R | F | 33 | 18 | 20 | 20/200 | C | Genetic / Developmental | 17 | R | M | 36 | 16 | 13 |
| 18 | R | M | 63 | 19 | 21 | 20/200 | C | Achromatopsia | 18 | R | M | 69 | 18 | 21 |
| 19 | R | F | 69 | 22 | 20 | 20/60 | A | Stargardt disease | 19 | R | F | 68 | 18 | 21 |
| 20 | R | F | 44 | 12 | 19 | 20/200 | A | Pseudotumor cerebri | 20 | R | F | 41 | 12 | 19 |
| 21 | R | M | 59 | 16 | 17 | 20/50 | A | Diabetic retinopathy | 21 | R | M | 62 | 16 | 17 |
| 22 | R | F | 38 | 16 | 20 | 20/100 | C | Genetic / Developmental | 22 | R | F | 42 | 18 | 21 |
| 23 | R | F | 34 | 18 | 22 | 20/300 | C | Rod cone dystrophy | 23 | R | F | 33 | 16 | 16 |
| - | 21R, 2B | 7M, 16F | 49.1 (17.3) | 17.4 (1.8) | 19.9 (1.3) | 20/195.0 (145.8) | 14C, 19A | - | - | 22R, 1B | 7M, 16F | 49.0 (16.3) | 16.1 (1.5) | 19.4 (2.0) |

**Figure 1.**
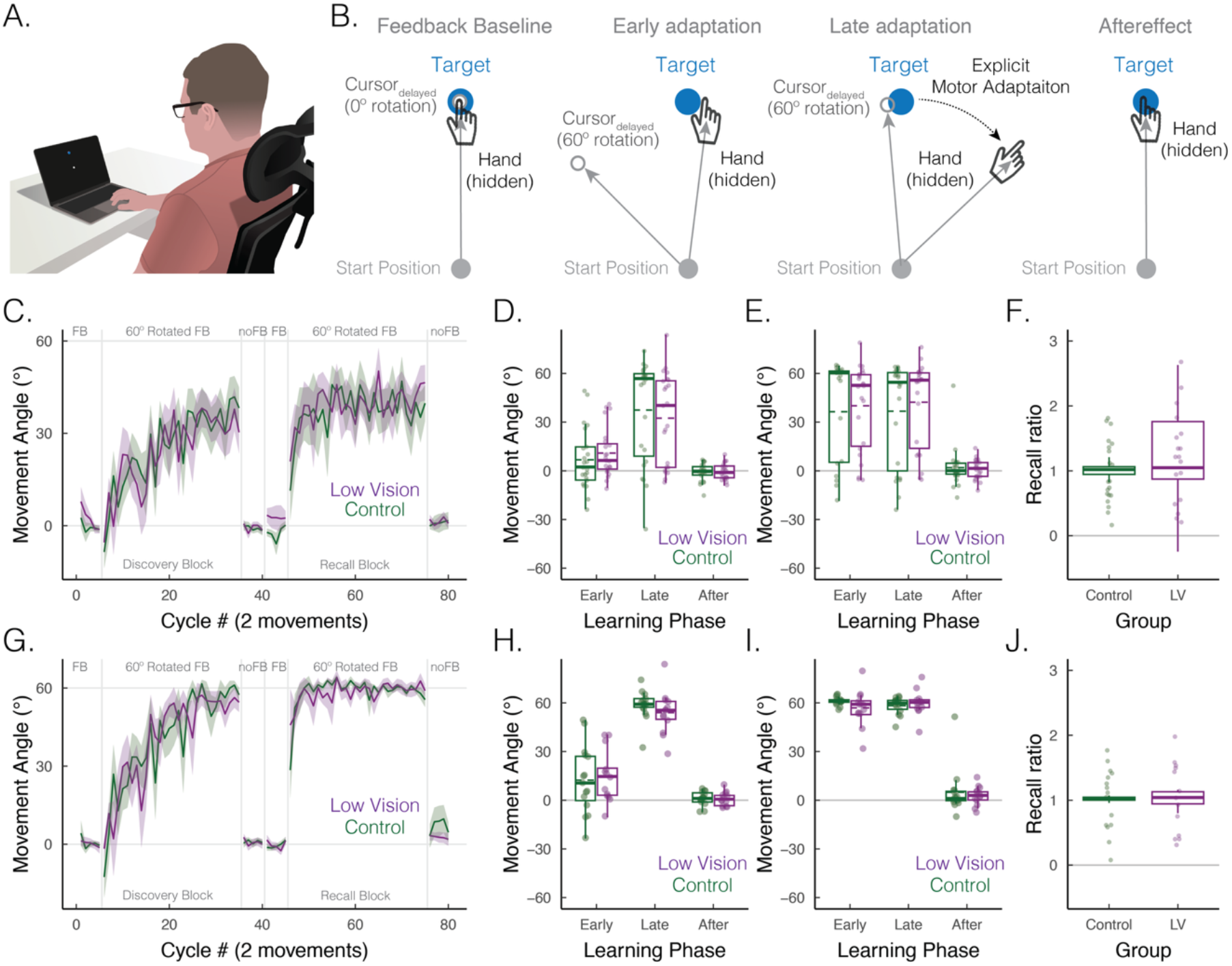
No association between low vision and impaired explicit motor adaptation. **(A)** Experiment setup. Participants used a trackpad to make rapid, goal-directed movements toward a visual target presented on a screen. **(B)** Delayed Feedback Task. After baseline trials with veridical feedback (cycles 1–10), participants completed a visuomotor rotation task with feedback rotated 60° clockwise or counterclockwise (counterbalanced across participants). Endpoint feedback was delayed by 800 ms—a manipulation known to suppress implicit adaptation and isolate explicit strategy use. Schematics illustrate hand and cursor positions during early adaptation (Discovery: cycles 7–16; Recall: cycles 47–56), late adaptation (Discovery: cycles 31–40; Recall: cycles 66–75), and the aftereffect phase (Discovery: cycles 41–45; Recall: cycles 76–80). Prior studies show that individuals typically do not look at their hand during such tasks, and feedback about their actual movement direction is withheld from view. **(C–F)** All participants. **(C)** Median hand angle (relative to the target at 0°) across cycles for Control (green) and Low Vision (purple) groups; shaded areas denote ±1 SEM. **(D)** Boxplots of hand angle during the Discovery block across Early, Late, and Aftereffect phases. (**E**) Same measures shown for the Recall block. **(F)** Recall ratio, calculated as Early Recall / Late Discovery hand angle, quantifying strategic recall. **(G–J)** Learners only (participants whose late adaptation significantly exceeded 0°). Same analyses as above. Boxplots show median (solid line), mean (dashed line), interquartile range, and full range. Individual dots reflect participant-level data.

To statistically evaluate performance, we focused on three a priori-defined phases: early adaptation, late adaptation, and aftereffect (Fig. 1C-E). We observed a significant main effect of phase (*F*(3, 308) = 24.6, *p* < 0.0001) and a significant interaction between phase and block (*F*(3, 308) = 8.4, *p* < 0.0001), indicating that movement angles varied across phases in a block-dependent manner: In the Discovery phase (Fig. 1D), participants exhibited modest early adaptation (8.9° [2.8°, 14.9°]; *t*(306) = 2.1, *p* = 0.14), followed by a substantial increase in late adaptation (34.9° [28.9°, 41.0°]; *t*(306) = 9.3, *p* < 0.001). Aftereffects were minimal (−0.7° [−6.8°, 5.3°]; *t*(308) = 0.48, *p* = 0.96), suggesting that learning was dominated by explicit strategies with little to no implicit adaptation. In the Recall phase (Fig. 1E), participants adapted rapidly from the outset. Early adaptation was robust (38.3° [32.2°, 44.3°]; *t*(306) = 10.4, *p* < 0.001) and remained high during late adaptation (39.5° [33.5°, 45.6°]; *t*(306) = 10.8, *p* < 0.001). As in the Discovery phase, aftereffects remained minimal (1.5° [−4.5°, 7.5°]; *t*(306) = 0.34, *p* = 0.99).

Post hoc comparisons confirmed significantly greater early adaptation in the Recall phase compared to Discovery (*t*(308) = 29.4, *p* < 0.001), reflecting the classic signature of “savings”—an accelerated re-expression of a previously learned strategy (Avraham et al., 2021; Hadjiosif et al., 2023; Morehead et al., 2015; Yin & Wei, 2020). However, late adaptation did not differ between blocks (*t*(308) = 1.3, *p* = 0.92), nor did aftereffects (*t*(308) = 0.6, *p* = 0.99). These findings suggest that although participants rapidly retrieved their explicit strategies during the recall block, overall learning remained unchanged, and implicit adaptation was not engaged in either case.

Turning to our main question, we asked whether low vision (LV) impacts the discovery or recall of an explicit strategy. We found no significant main effect of group (*F*(1, 222) = 0.2, *p* = 0.69), nor any significant interactions involving group (all *p*s > 0.62), indicating that performance did not differ between LV and Control participants across any phase or block. A complementary Bayesian analysis yielded a Bayes Factor of 5.6 in favor of the null, providing substantial evidence *against* a group difference. As shown in Fig. 1C, the LV group’s learning curves closely mirrored that of the Control group.

Recognizing that movement angle alone may not fully capture recall—given variability in strategy use during the initial exposure—we conducted a complementary analysis. Specifically, we computed a “recall ratio,” defined as the movement angle during early adaptation in the Recall phase normalized by the angle during late adaptation in the Discovery phase. A ratio of 1 indicates complete recall of the previously learned strategy, values below 1 reflect partial recall, and values above 1 suggest the use of a larger aiming strategy upon re-exposure to the perturbation.

Using this measure, both groups demonstrated strong recall (Fig. 1F). The Control group had a median recall ratio of 1.0 (IQR: 0.9–1.1; Wilcoxon test, Recall Ratio vs. 0: *V* = 251, *p* < 0.001), and the LV group also had a median recall ratio of 1.0 (IQR: 0.9–1.8; *V* = 235, *p* = 0.002). Importantly, the recall ratio did not differ significantly between groups (*W* = 307, *p* = 0.36), providing further evidence that strategic recall remains intact in individuals with low vision.

As shown in Fig. 1D–F, participants showed substantial variability in both the discovery and recall of an explicit aiming strategy. Consistent with prior findings, a subset of individuals failed to adapt to the perturbation by the end of the Discovery block—that is, their late adaptation angles did not differ significantly from baseline. This included 8 participants in the Control group (35%) and 10 in the Low Vision group (43%), with comparable proportions across groups (Fisher’s exact test, p = 0.76, odds ratio = 0.70). These “non-learners” may represent a broader class of participants who either did not attend to the feedback, misunderstood the task instructions, or attempted to adapt but failed to discover an effective strategy. As a result, their data may obscure potential group differences in strategy use, and their recall ratios are uninterpretable, as no strategy was established to recall.

We thus performed a secondary analysis limited to “learners,” defined as participants whose movement angle during the late adaptation epoch of the Discovery block was significantly greater than zero. This subset included 15 participants from the Control group and 13 from the LV group: Our key findings were replicated (Fig. 1G-J). There was no main effect of group (*F*(1, 202) = 0.01, *p* = 0.92) and no significant interactions involving group (all *p*s > 0.36), across all phases. As before, both groups showed strong recall (IQR, Control: 1.0–1.0; LV: 0.9–1.1), with no significant difference between them (*W* = 101, *p* = 0.89), reaffirming that low vision had minimal impact on the ability to discover and recall an explicit aiming strategy.

## Discussion

Low vision is known to impair the ability to discriminate the spatial position of visual objects (Massof & Fletcher, 2001; Timmis & Pardhan, 2012), leading to slower and less accurate goal-directed movements (Cheong et al., 2021; Endo et al., 2016; Jacko et al., 2000; Kotecha et al., 2009; Lenoble et al., 2019; Timmis & Pardhan, 2012; Verghese et al., 2016). However, the impact of visual uncertainty from low vision on motor adaptation—particularly explicit strategy use—remains poorly understood. To address this, we used a visuomotor adaptation task specifically designed to isolate explicit re-aiming. Strikingly, low vision had no measurable effect on strategy use, during either initial discovery or later recall. Despite visual impairments, individuals with low vision were just as effective as controls in forming and retrieving a deliberate, compensatory strategy.

These findings challenge the assumption that degraded visual input uniformly impairs all forms of visuomotor learning. While prior work has shown that visual uncertainty attenuates implicit adaptation—particularly when feedback is noisy or imprecise—our results suggest that explicit learning mechanisms may be more resilient. This dissociation between the effects of visual uncertainty on implicit and explicit processes adds to a growing literature suggesting distinct computational and neural underpinnings for these learning systems (Areshenkoff et al., 2023; Cisneros et al., 2024; Kim et al., 2020).

Several factors may explain why explicit strategy use remained intact in individuals with low vision. One possibility is that chronic visual uncertainty engages compensatory mechanisms that preserve strategic control—similar to what we previously observed in individuals with proprioceptive deafferentation (Tsay, Chandy, et al., 2024). Specifically, individuals with low vision may learn to rely more heavily on intact regions of their visual field (Cloutier & DeLucia, 2022; Crossland et al., 2011) or develop heightened neural sensitivity to visual errors over time (He et al., 2006). Alternatively, intact strategy use may reflect task conditions in which visual errors were large, unambiguous, and well above perceptual thresholds. In more subtle conditions—where errors are smaller or lower in contrast (Casasnovas et al., 2024)—strategy discovery in individuals with low vision may suffer.

Clinically, the finding that strategy use remains largely intact in low vision—and more broadly, resilient to visual uncertainty—has important implications. Explicit strategies are central to many rehabilitation protocols, where patients with movement disorders are taught to consciously compensate for changes in body state (e.g., muscle weakness) or environment (e.g., prosthetic use) following neurological injury (Abram et al., 2025; Leech et al., 2021; Roemmich & Bastian, 2018; Tsay & Winstein, 2020). Our results suggest that even individuals with impaired vision can benefit from strategy-based interventions. Rehabilitation specialists may leverage this preserved capacity by providing clear, goal-oriented instructions and emphasizing task-relevant errors, thereby promoting effective strategic compensation when sensory feedback is limited.

Nonetheless, several limitations warrant consideration. First, the absence of group differences may reflect limited severity; more profound visual impairments may be necessary to disrupt strategy use. However, our sample spanned a broad range of diagnoses and functional impairments, capturing the diversity of individuals affected by low vision. Second, visual function was assessed using relatively coarse clinical criteria—specifically, records indicating uncorrectable visual acuity worse than 20/50. Without more fine-grained measures (e.g., contrast sensitivity, visual field mapping, psychophysical thresholds, or detailed diagnostic classifications), we cannot fully characterize the extent of impairment or individual variability within the low vision group. Third, the experiments were conducted remotely, limiting our ability to fully control and standardize the testing environment. This approach was essential for reaching individuals who could not easily travel to the lab and allowed data collection to continue during the COVID-19 pandemic, when the study was conducted (Fooken et al., 2023; Tsay, Steadman, et al., 2024). To ensure data quality, we implemented several safeguards: Participation was restricted to trackpad users to standardize input devices. We focused on group-level comparisons, where individual variability (e.g., screen size, refresh rate) likely averages out. Real-time phone support from the experimenter further ensured consistency across participants. Notably, this remote format may be especially advantageous, as neither group benefited from in-person interaction with the experimenter. Finally, our results replicated hallmark features of explicit strategy use—such as minimal aftereffects— commonly observed in lab-based settings, supporting the validity of our online approach.

Together, these findings suggest that explicit mechanisms underlying sensorimotor adaptation are resilient to visual uncertainty in individuals with low vision. This insight highlights the potential to leverage explicit strategies in motor rehabilitation, even when visual input is degraded.

## Methods

### Participants

All participants gave written informed consent in accordance with policies approved by the UC Berkeley Institutional Review Board. Participation in the study was in exchange for monetary compensation. We recruited 23 individuals with low vision. Our sole inclusion criterion was visual acuity worse than 20/50 (0.4 logMAR) in the better-seeing eye, uncorrectable with refractive aids (i.e., glasses or contact lenses), as confirmed by clinical records. Participants were recruited through UC Berkeley’s Meredith W. Morgan University Eye Center and a registry maintained by the National Research & Training Center on Blindness and Low Vision at Mississippi State University (Table 1). Those who consented completed a phone-based screening, including a medical history review and the Montreal Cognitive Assessment–Blind (MoCA-Blind) to assess general cognitive function (28). In the low vision group, visual acuity in the better-seeing eye had a mean of 20/195.0 (SD = 145.8). Of these participants, 13 had congenital visual impairments and 10 had acquired impairments.

We also recruited 23 age-matched controls from the UC Berkeley community (Table 1). All control participants had normal or corrected-to-normal vision. As intended, the low vision and control groups did not differ significantly in age (t(44) = 0.06, p = 0.95), handedness (χ^2^(1) = 0, p = 1), or sex (χ^2^(1) = 0, p = 1). Although the low vision group had, on average, one year less education (t(44) = 2.7, p = 0.01), cognitive ability—as measured by the MoCA-Blind—did not differ between groups (t(37) = 1.0, p = 0.33).

### Apparatus

Participants used their own laptop or desktop to access a custom webpage hosting the experiment (Tsay, Lee, et al., 2021; Tsay et al., 2020) and performed the motor learning task using a trackpad (sampling rate ∼60 Hz) (Fig. 1A). Stimulus size and position were scaled to each participant’s screen dimensions. For reference, the parameters reported below correspond to a 13” diagonal monitor. Throughout the one-hour session, the experimenter remained on the phone to provide instructions and support.

### General Procedures

Participants performed reaching movements across a virtual workspace on their screen. Each trial involved a movement from the center of the workspace (white circle, 0.5 cm diameter) to a visual target (blue circle, 0.5 cm diameter). The radial distance from the center to the target was 6 cm. Targets appeared at one of two locations: 60° (upper right) or 210° (lower left).

Each trial began when the participant moved their cursor into the starting circle. After the cursor remained in the start circle for 500 ms, the blue target appeared. Participants were instructed to make a rapid, slicing movement through the target following an auditory go-cue (a single beep). A delay of 1200 ms (±100 ms jitter) was imposed between target appearance and the auditory go-cue to standardize movement preparation time across participants and minimize speed–accuracy trade-offs. If participants moved before the go-cue, they heard the message “Wait for the tone!” If they failed to initiate movement within 800 ms after the cue, they heard “Move earlier!”

The cursor disappeared as soon as the hand left the start circle. Visual feedback during center-out movements took one of three forms: veridical, rotated, or no feedback. In veridical feedback trials, the cursor appeared at the movement endpoint and accurately reflected the hand’s direction (Fig. 1B). In rotated feedback trials, the cursor was shown at the endpoint but offset by 60° from the actual movement angle once the movement distance exceeded 6 cm—the target distance. The direction of rotation (clockwise or counterclockwise) was counterbalanced across participants. In no-feedback trials, the cursor remained off throughout the movement. To prevent feedback from influencing the return movement to start the next trial, the cursor was only visible within 2 cm of the start location. A single trial consisted of one complete outward movement from the start circle to a target. A movement cycle comprised two center-out reaches—one to each target.

### Experimental Design

First, participants watched an instructional video introducing key task features, including the go-cue and delayed cursor feedback. Second, they completed 15 practice trials to familiarize themselves with the web-based reaching environment and both veridical and no-feedback conditions. Third, participants completed the main task consisting of six blocks separated into two phases: initial exposure to a perturbation (assessing Discovery) and re-exposure (assessing Recall). Impaired discovery would manifest as reduced performance in the first phase, while impaired recall would emerge in the second. This design allows us to test how visual uncertainty influences the strategic processes that support sensorimotor learning. In the Discovery phase, they were first introduced to the adaptation paradigm in three blocks: baseline with veridical feedback (5 cycles), rotated feedback for strategy discovery (30 cycles), no-feedback aftereffect block (5 cycles). This phase was immediately followed by an identical Recall phase comprising the same three blocks. Across both phases, this totaled to 80 movement cycles across two targets = 160 trials total.

Before each veridical feedback block, participants were instructed: “Please move your white cursor directly to the blue target immediately after the tone.” Before each rotated feedback block, they were told: “Your white cursor will appear at an offset from your movement. To hit the blue target with your white cursor, please reach somewhere different than the blue target immediately after the tone.” Prior to the no-feedback blocks, the instruction was: “Your white cursor will be hidden and no longer offset from where you moved. Please move directly and immediately to the blue target after the tone.” The experimenter remained on the phone and assessed comprehension by prompting participants to restate the instructions in their own words. No participants were excluded based on this understanding check.

### Data Analysis

All data and statistical analyses will be performed in R. The primary dependent variable was the endpoint movement angle on each trial (i.e., the angle of the cursor when the movement amplitude reached a 6 cm radial distance from the start position relative to the location of the target). Because no target-specific differences were expected, movement angles were averaged across the two target locations to simplify visualization.

We compared movement angle between groups in three a priori defined epochs (Tsay, Schuck, et al., 2022): Early adaptation, late adaptation, and aftereffect. These epochs were examined separately in the Discovery and Recall phases. Early adaptation was defined as the initial ten movement cycles after the rotation was introduced (Discovery: cycles 7 – 16; Recall: cycles 47 – 56). Late adaptation was defined as the final ten movement cycles of the rotation blocks (Discovery: cycles 31 – 40; Recall: cycles 66 – 75). Aftereffect was defined as all movement cycles without visual feedback after the rotation was removed (following Discovery: cycles 41 – 45; following Recall: cycles 76 – 80).

We used F-tests with the Satterthwaite approximation to assess the significance of coefficients (beta values) from linear mixed-effects models (R functions: lmer, lmerTest, ANOVA, emmeans). Pairwise post hoc comparisons were conducted using two-tailed t-tests or Wilcoxon signed-rank tests when parametric assumptions were violated. P-values were adjusted for multiple comparisons using the Tukey method. When variances between groups were unequal, degrees of freedom were adjusted accordingly.

## Acknowledgements

We thank Matthew Warburton for illustrating our experimental setup.

